# Molecular Profiling to Infer Neuronal Cell Identity: Lessons from small ganglia of the Crab *Cancer borealis*

**DOI:** 10.1101/690388

**Authors:** Adam J. Northcutt, Daniel R. Kick, Adriane G. Otopalik, Benjamin M. Goetz, Rayna M. Harris, Joseph M. Santin, Hans A. Hofmann, Eve Marder, David J. Schulz

**Affiliations:** Division of Biological Sciences, University of Missouri-Columbia, Columbia, MO USA 65211; Neural Systems and Behavior Course, Marine Biological Laboratory, Woods Hole, MA, USA 02543; Volen Center and Biology Department, Brandeis University, Waltham, MA USA 02454; Center for Computational Biology and Bioinformatics, The University of Texas at Austin, Austin, TX, USA 78712; Department of Integrative Biology and Institute for Cellular and Molecular Biology, The University of Texas at Austin, Austin, TX, USA 78712; Institute for Neuroscience, The University of Texas at Austin, Austin, TX, USA 78712

**Keywords:** qPCR, RNA-seq, ion channel genes, receptor genes

## Abstract

Understanding circuit organization depends on identification of cell types. Recent advances in transcriptional profiling methods have enabled classification of cell types by their gene expression. While exceptionally powerful and high throughput, the ground-truth validation of these methods is difficult: if cell type is unknown, how does one assess whether a given analysis accurately captures neuronal identity? To shed light on the capabilities and limitations of solely using transcriptional profiling for cell type classification, we performed two forms of transcriptional profiling – RNA-seq and quantitative RT-PCR, in single, unambiguously identified neurons from two small crustacean networks: the stomatogastric and cardiac ganglia. We then combined our knowledge of cell type with unbiased clustering analyses and supervised machine learning to determine how accurately functionally-defined neuron types can be classified by expression profile alone. Our results demonstrate that expression profile is able to capture neuronal identity most accurately when combined with multimodal information that allows for post-hoc grouping so analysis can proceed from a supervised perspective. Solely unsupervised clustering can lead to misidentification and an inability to distinguish between two or more cell types. Therefore, our study supports the general utility of cell identification by transcriptional profiling, but adds a caution: it is difficult or impossible to know under what conditions transcriptional profiling alone is capable of assigning cell identity. Only by combining multiple modalities of information such as physiology, morphology or innervation target can neuronal identity be unambiguously determined.

**SIGNIFICANCE STATEMENT:** Single cell transcriptional profiling has become a widespread tool in cell identification, particularly in the nervous system, based on the notion that genomic information determines cell identity. However, many cell type classification studies are unconstrained by other cellular attributes (e.g., morphology, physiology). Here, we systematically test how accurately transcriptional profiling can assign cell identity to well-studied anatomically- and functionally-identified neurons in two small neuronal networks. While these neurons clearly possess distinct patterns of gene expression across cell types, their expression profiles are not sufficient to unambiguously confirm their identity. We suggest that true cell identity can only be determined by combining gene expression data with other cellular attributes such as innervation pattern, morphology, or physiology.

## INTRODUCTION

Unambiguous classification of neuronal cell types is a long-standing goal in neuroscience with the aim to understand the functional components of the nervous system that give rise to circuits and, ultimately, behavior (1–6). Beyond that, agreement upon neuronal cell types provides the opportunity to greatly increase reproducibility across investigations, allows for evolutionary comparisons across species (7, 8), and facilitates functional access to and tracking of neuron types through developmental stages (9). To this end, attempts at defining neuronal identity have been carried out using morphology, electrophysiology, gene expression, spatial patterning, and neurotransmitter phenotypes (10–18). Since the earliest efforts to capture the transcriptomes of single neurons, using linear or PCR amplification of mRNA followed by either cDNA library construction (19) or microarray hybridization (10, 20, 21), scRNA-seq (22) has become the method of choice for many genome-scale investigations into neuron cell type. Advances in microfluidics, library preparation, and sequencing technologies have propelled an explosion of molecular profiling studies seeking to use unique gene expression patterns to discriminate neuronal types from one another, whether for discovery of new types or further classification of existing ones (23, 24, 33–36, 25–32).

Molecular profiling approaches to tackle the problem of neuronal cell identity have many advantages: first, single-cell transcriptomic data contain thousands of measurements in the form of gene products that can be used both in a qualitative (in the form of marker genes) and quantitative (in the form of absolute transcript counts) manner (6). Second, scRNA-seq allows for very high-throughput processing of samples with hundreds, if not thousands, of single cell transcripts simultaneously using barcoding techniques (37). Third, these techniques can be applied to species that lack well-annotated transcriptomic information, as the cost to generate *de novo* reference transcriptomes has decreased dramatically in recent years (38). Even the sequencing of heterogeneous tissues from the central nervous system (CNS) can be used in conjunction with predictive modeling to reconstruct markers for major classes of CNS cell types, as has been done with oligodendrocytes, astrocytes, microglia, and neurons, in both humans and mice (39). Classifying neurons into different major categories (such as excitatory vs inhibitory, parvalbumin^+^ vs parvalbumin^-^, etc.) using qualitative expression measures is an easier task than quantitative approaches that separate neurons into smaller subclasses, but runs into limitations as to how far further classification can proceed. Subclasses of neuron types likely require greater depth of sequencing to resolve, and these neurons are more likely to be defined by the expression of multiple genes rather than unique markers (40). Yet this also is an inherent limitation of scRNA-seq: low abundance transcripts are often missed or inaccurately classified as differentially expressed (41), and methods to dissociate and isolate cells can alter their transcriptomic profiles before they are even measured (42, 43).

There now have been many studies seeking to determine how many transcriptomically-defined cell types might be present in a given part of the brain. For instance, an initial study of the cell type diversity of the mouse primary visual cortex revealed 42 neuronal and 7 non-neuronal cell types (25). More recent work from the same group identified 133 transcriptomic cell types (44). Work in the retina has led the way as an example of generating a cell type consensus with an unknown endpoint. Multimodal information of retinal ganglion cell properties, including morphology, physiology, gene expression, and spatial patterning, has converged on over 65 cell types in the macaque fovea and peripheral retina (45). However, not all systems have the same technical advantages as the retinal ganglion cells (such as uniform spatial patterning) that can be indicative of cell type, and multimodal information can be more difficult to obtain than high-throughput transcriptomic profiling methods. Therefore, the reliability of transcriptomic profiling with respect to neuronal identity requires additional evaluation.

In this study, we validate and compare transcriptional profiling via scRNA-seq and qRT-PCR methods, using supervised and unsupervised analyses, in model systems in which neurons are unambiguously identified based on electrophysiological output, synaptic connectivity, axonal projection, and innervation target: the stomatogastric (STG) and cardiac ganglia (CG) of the crab, *Cancer borealis*. This approach allows us to directly test how much of the known functional identity of a neuron is captured in the transcriptomic profile of single neurons within a given network.

## RESULTS

### Molecular Profiling of Single Identified STG and CG Neurons by RNA-seq

Because of their large individual cell body size and our abiity to manually collect single identified STG neurons (Fig. 1), we generated transcriptomes for Pyloric Dilator (PD; N=11), Gastric Mill (GM; N=11), Lateral Pyloric (LP; N=8), and Ventricular Dilator (VD; N=8) neurons by typical library preparations rather than more automated procedures such as Drop-seq, Split-Seq, or 10X Genomics (46). Sequencing data were mapped to the *C. borealis* nervous system transcriptome (47). After removing transcripts for which there was no expression in any cell type, our data set contained 28,459 distinct contigs (i.e. contiguous sequences) in the complete RNA-seq data set. These contigs represent more than the full set of genes transcribed in these cells, as multiple contigs may map to a single gene but during transcriptome assembly the intervening sequence could not be resolved to assemble these distinct fragments (see [58]). We then began our analysis of these data using unbiased hierarchical clustering methods, as is commonly done in this field. Using the complete data set (referred to as “All Expressed Contigs”), hierarchical clustering (with data centered and scaled across contigs) resulted in five clusters (Fig. 2A) that appeared not to segregate by cell type. One exception was observed among PD cells. All but two PD cells fell within one distinct cluster, albeit with a GM cell also identified in this cluster (Fig. 2A). While not surprising, the complete cellular transcriptome on its own does not distinguish cell types.

**Figure 1.**
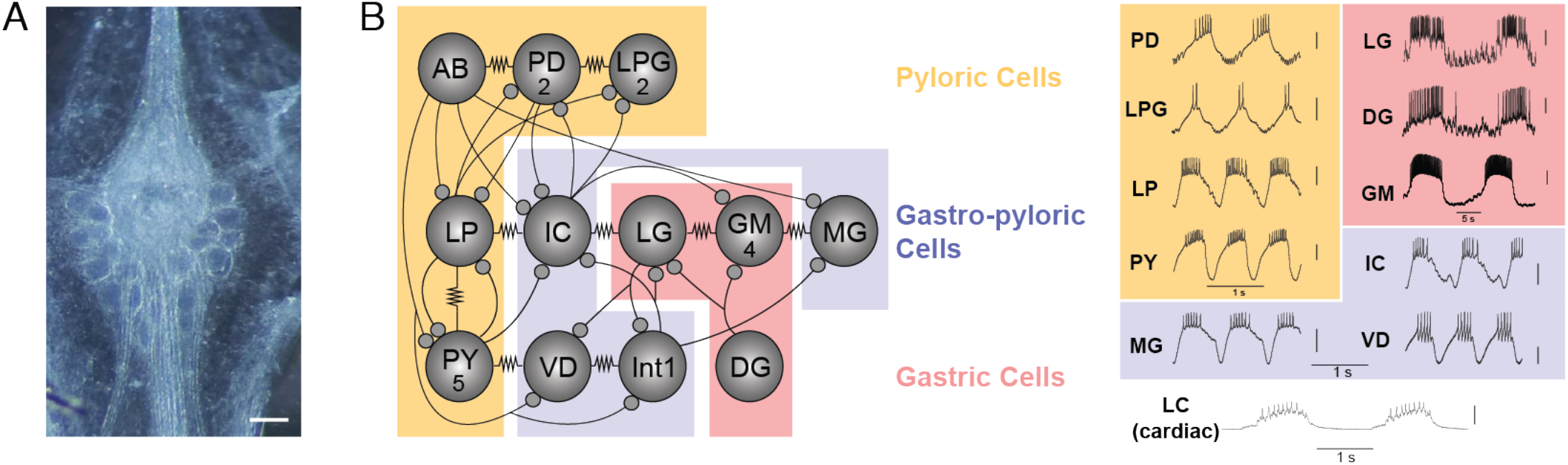
**A)** Photomicrograph of the stomatogastric ganglion. Scale bar = 200 μm. **B)** Circuit map of the stomatogastric ganglion (STG). The STG contains 12 cell types that innervate the pylorus and gastric mill of the crab stomach. These cells are individually identifiable, and their chemical (closed circles) and electrical (resistor symbols) synaptic connections are all known. We used 10 of these 12 STG cell types (not AB or INT1) for this study, as well as motor neurons of the cardiac ganglion as an outgroup for comparison. Example traces taken from intracellular recordings of each the 11 identified neuron types used in this study. Neurons are involved in three different networks/circuits in the crab, *Cancer borealis*: the pyloric network (PD, LPG, LP, and PY; orange box), the gastric network (LG, DG, and GM; red box) and the cardiac ganglion network (bottom). Note the time scale difference in the long-lasting bursts of the gastric cells (red box) relative to the pyloric cells (orange box). Some neurons (IC, VD, and MG) participate in both gastric and pyloric network activity, and are noted in the purple box. Large Cell (LC) motor neurons of the cardiac ganglion are used as an “outgroup” to compare expression patterns of motor neurons from a distinct ganglion (cardiac ganglion). Each of the representative recordings is independent as an example of individual cell output, and simultaneous network activity is not plotted here. Thus, none of the phase relationships of these units within their respective rhythms is implied in any of the recordings.

**Figure 2.**
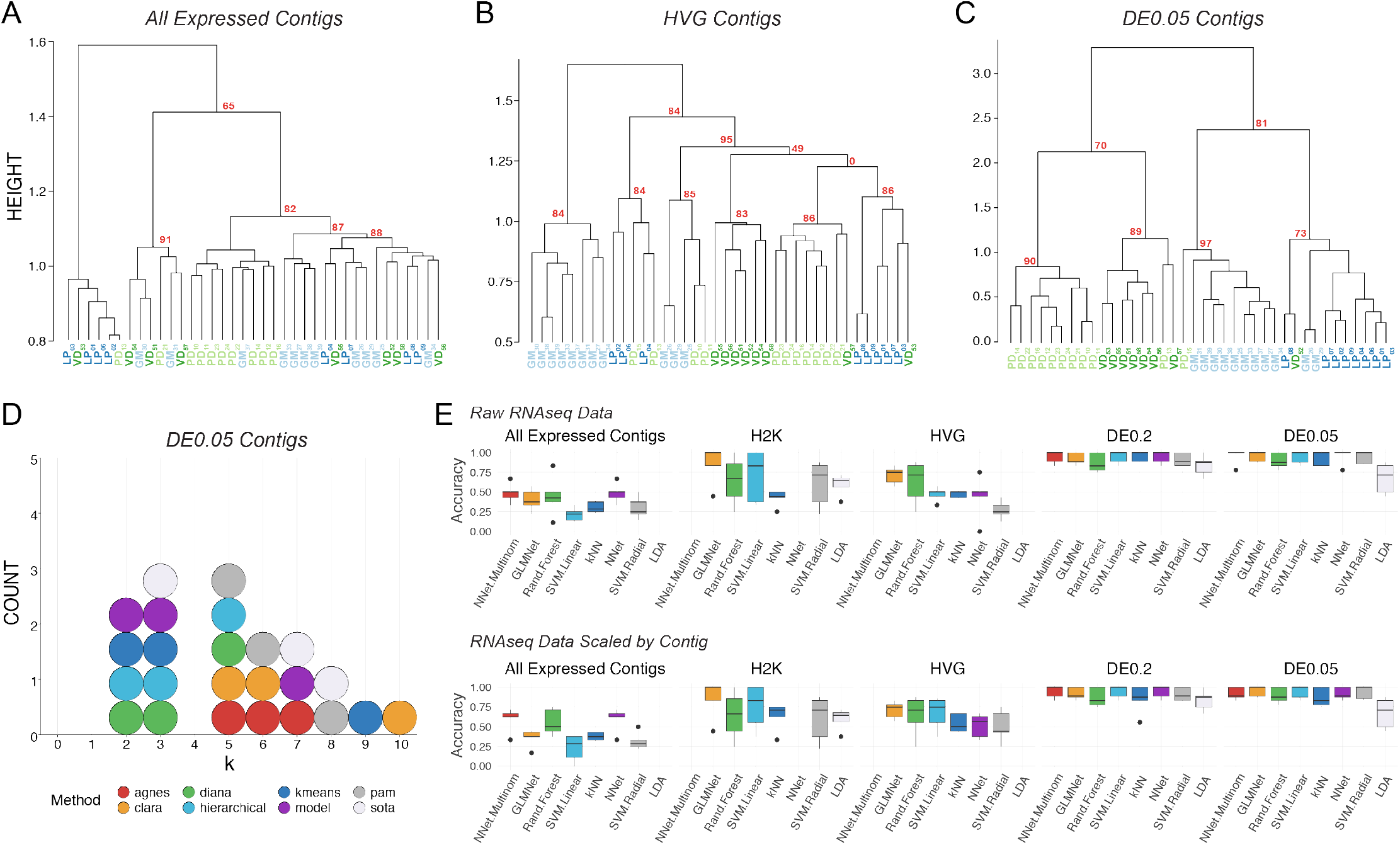
Post-hoc recapitulation of cell identity via single cell RNAseq with hierarchical clustering and supervised machine learning (sML) algorithms. **A)** Hierarchical clustering of cell type with correlation as the distance metric, Ward.D2 as the clustering method, and data centered and scaled by contig for all expressed contigs, **B)** Highly Variable Gene (HVG) dataset, and **C)** differentially expressed (DE) contigs at the q < 0.05 level. Each cell type is color coded, and Approximately Unbiased (AU) p-values are noted for each of the major nodes. Cells are identified by type (LP, PD, GM, VD) and a subscript that denotes a unique sample identifier. **D)** Dotplot of the top 3 predicted number of clusters (k values) for eight algorithms. None of these algorithms correctly predicted the expected 4 distinct clusters that would represent the 4 different cell types in this assay. **E)** Accuracy (proportion of correctly identified cells) of cell type prediction using 8 different methods of sML (generalized linear model (GLM), k-Nearest Neighbors (kNN), Neural Network (NN), Multinomial Neural Network (MNN), Random Forest (RF), Support Vector Machine with a linear kernel (SVML), Support Vector Machine with a radial kernel (SVMR), and Linear Discriminant Analysis (LDA)) for each of the data sets. Box and whisker plots show the efficacy of these methods to recapitulate cell identity from these two sets of contigs as estimated by cross validation (5 folds). To assess the efficacy of these methods on the full RNA seq dataset, we used principle component analysis (PCA) for dimensionality reduction (i.e. >28,000 contigs to 38 PCs) while retaining 99% of the variance. Results are shown for raw data (*top row*) and data scaled across contigs (*bottom row*).

We identified and extended our unbiased analysis to the most variably expressed genes in the RNA-seq dataset. The first subset represents the top 2000 most variable contigs (referred to as the “H2K contigs”) and the second subset includes variable genes identified using a method described by Brennecke et al. (48), assuming a false discovery rate of 0.2, which resulted in 922 contigs (referred to as “HVG contigs”). Focusing on variably expressed contigs improved clustering with respect to cell identity, with the HVG dataset outperforming the H2K. In the HVG clustering (Fig. 2B), 8/11 GM cells, 5/8 VD cells, 5/8 PD cells, and 5/8 LP cells formed distinct clusters. However, these nodes are not perfectly segregated by cell type and cells of each kind fail to appropriately cluster. If blind to these cell types, the HVG clustering analysis yields 5-6 distinct cell-type clusters, rather than the appropriate 4 (Fig. 2B).

To achieve the best performance possible with scRNA-Seq clustering analyses, we unblinded the analyses to cell type and selected only differentially expressed transcripts. We selected two pools of differentially expressed transcripts: those with a 2-fold or higher level of expression difference and a q-value < 0.2 (referred to a “DE0.2”) or q-value <0.05 (“DE0.05”). Of course, differential expression (DE) analysis can only be carried with *a priori* knowledge of cell identity or some other post-hoc feature by which samples can be grouped. DE analysis with a q-value cutoff of 0.2 identified 137 transcripts (DE0.2), while a q-value of 0.05 identified only 45 transcripts (DE0.05). Hierarchical clustering of the q<0.2 data set resulted in better clustering, but still failed to faithfully recapitulate cell identity. Hierarchical clustering was greatly improved by using the q<0.05 dataset (DE0.05; Fig. 2C) but remained imperfect.

To reveal which preprocessing and clustering methods best recapitulate the predicted number of clusters based on known cell identity, we applied eight cluster estimation algorithms (optCluster package (49)) on the DE0.05 data set (centered and scaled by contig, Ward.D2 and a correlation dissimilarity matrix; Fig. 2D). The highest performing clusterings using the DE0.05 data resulted from using Ward’s D with a correlation distance metric, resulting in a Jaccard index of 0.738. The results of cluster estimation differed based on the preprocessing of the datasets. Cluster estimation algorithms were selected from a set of 10 algorithms for use with continuous data as they all yielded usable output. We retained the top three predicted k values from each. When data were centered and scaled by contig (Fig. 2D), the mode number of clusters estimated was 3 (5 indices) and 5 (5 indices), and none predicted the correct number of 4 clusters.

Finally, to assess whether unblinded analyses could predict cell type, we tested the ability of 8 supervised machine learning (sML) classification algorithms (generalized linear model (GLM), k-Nearest Neighbors (kNN), Neural Network (NN), Multinomial Neural Network (MNN), Random Forest (RF), Support Vector Machine with a linear kernel (SVML), Support Vector Machine with a radial kernel (SVMR), and Linear Discriminant Analysis (LDA)) to sort cells based on their transformed or untransformed mRNA abundances. Each model’s accuracy on new data was estimated using 5-fold cross validation. To capture the variation in the All Expressed Contigs dataset, we transformed the data with PCA and used the first 38 principal components, which accounted for over 99% of the variation. The sML mean accuracies on the All Expressed Contigs (PCA transformed) data set were extremely low, with a maximum mean accuracy of 48.6% (Fig. 2E). sML accuracies improved substantially when classifying the RNA-seq data preprocessed to identify variably expressed contigs (H2K, HVG) and DE contigs (DE0.2, DE0.05), often producing 100% accuracy for several cross-validation folds (Fig. 2E). It should be noted that no method classified all folds with complete accuracy, even with only DE contigs– most methods ranged between 75% to 100% accuracy. While these results are encouraging, even under optimal conditions (transcriptomic data, selection of transcripts by differential expression, ability to use supervised methods) we were unable to consistently classify these neurons with 100% accuracy.

### Principal Component Analysis of scRNA-seq Datasets

Principal Component Analysis (PCA) is often used to determine if the variance seen among transcript abundances can be used to separate cells into discrete types. Thus, we performed PCA on the four RNA-seq datasets (H2K, HVG, DE0.2, DE0.05) to examine the ability of this approach to discriminate among cell types (Fig. 3). For most of these datasets, the first principal component (PC1) accounted for >40% of the explained variance, with the exception of the HVG dataset (Fig. 3). As such, we have listed the top 10 contigs contributing to variation in PC1 for all four datasets in Table S1. We generated pairwise plots of all three PCs in attempts to visualize separation of samples into distinct cell types. There is little ability to resolve cell type differences in the H2K and HVG datasets (Fig. 3A, 3B). However, the differentially expressed transcripts allow for some separation of cell type (Fig. 3C, 3D), with PD becoming somewhat distinct for example in the DE0.05 dataset (Fig. 3D).

**Figure 3.**
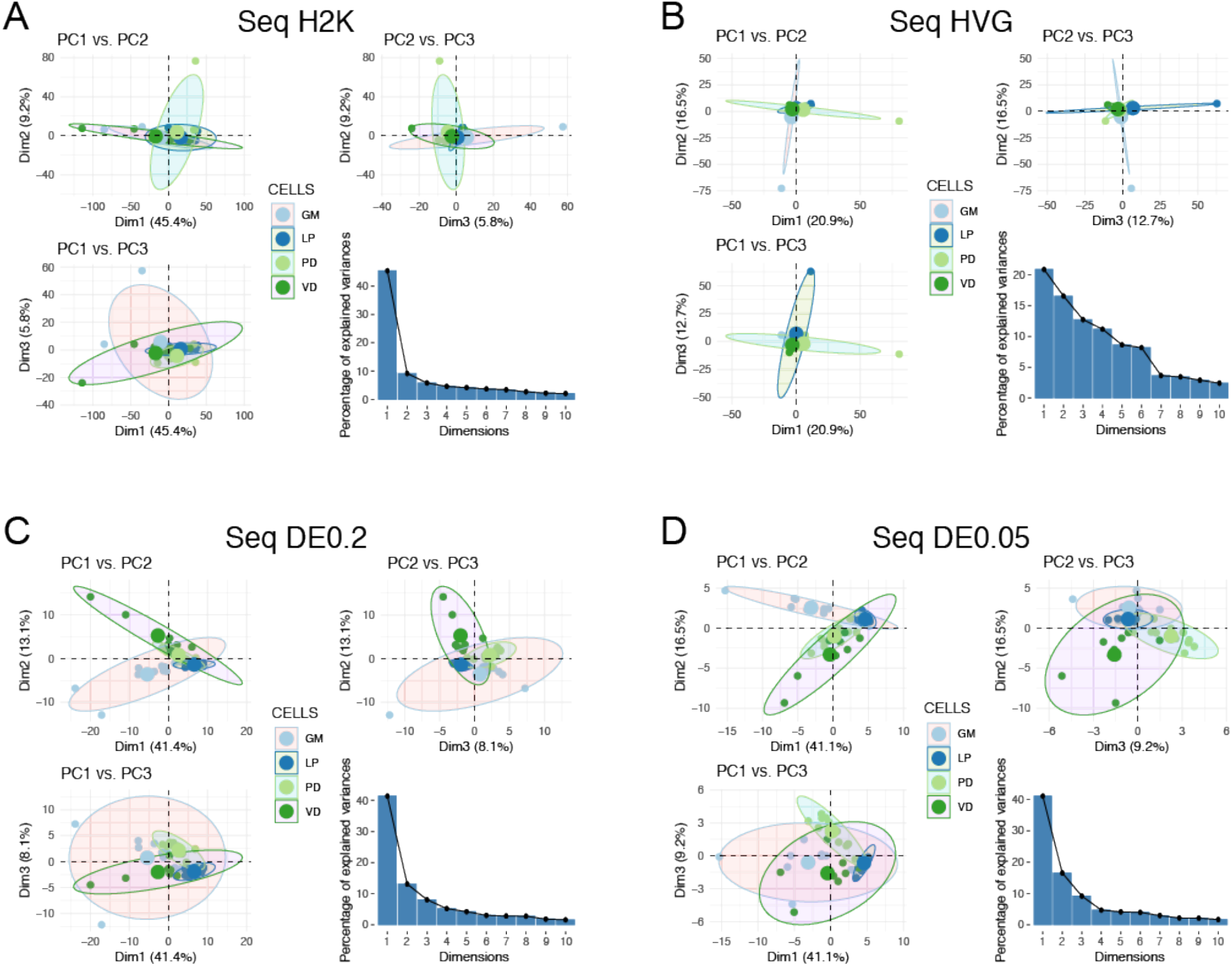
Principal Component Analysis (PCA) for four different RNAseq datasets. We performed PCA using **A)** the 2,000 contigs with the highest variance in expression (H2K), **B)** the Highly Variable Gene set (HVG), and differentially expressed (DE) contigs at the **C)** q < 0.2 (DE0.2) and **D)** q < 0.05 (DE0.05) levels. For each panel we have plotted pairwise comparisons of PC1, PC2, and PC3, as well as a scree plot representing the percentage of variance explained by PCs 1-10.

### Gene Ontology Analyses of RNA-seq Datasets

To determine the types of genes represented in our most variable (H2K and HVG) and differentially expressed (DE0.2, DE0.05) data sets among cell populations, we performed Gene Ontology (GO) Enrichment Analysis using analysis tools from the PANTHER Classification System (50). Because there is relatively little gene annotation work in the crab, we performed GO analysis by first using BLAST to find the top *Drosophila* ortholog for a given contig, and then retrieving the GO terms associated with this ortholog for analysis. Thus while this analysis provides interesting insight into cell-type specific differences in gene expression, there are limitations to the interpretation, particularly with regards to fold enrichment in *Drosophila* relative to crab. The most robust expression differences (highest Fold Enrichment) in the H2K Molecular Function dataset were those of ATP-synthase activity and clathrin binding (Table S2). Others of note include mRNA-3’UTR binding, cell adhesion molecule, and calcium ion binding (Table S2). More resolution is gained by examining the Biological Process category, where H2K contigs were most overrepresented for “regulation of short-term neuronal synaptic plasticity,” “positive regulation of neuron remodeling,” “substrate adhesion-dependent cell spreading,” and “clathrin-dependent synaptic vesicle endocytosis” categories (Table S3) among many others. The HVG dataset shows relatively few enriched categories (Tables S4 and S5) with FDR correction employed, including ATP binding and transferase activity (related to acetylcholine synthesis).

The differentially expressed contigs of the DE0.2 data set showed no significantly enriched contigs with FDR employed. Without any p-value correction, a number of molecular function categories appear as enriched (Table S6). However, this is less an appropriate enrichment analysis (due to the relatively small number of contigs) and more a description of gene categories present in the DE0.2 contigs. The top several hits are all indicative of transmitter phenotype, particularly acetylcholine synthesis (Table S6). However, other receptor activity is represented, such as GABA-gated chloride channel and GABA-A receptor activity. Finally, cell-cell adhesion mediator activity appears once again in this list.

### Molecular Profiling of Single Identified STG and CG Neurons Using Candidate Genes

One class of genes that we were surprised to not see represented in DE analyses were the voltage-gated ion channels. A very recent study found that three classes of neuronal effector genes - ion channels, receptors and cell adhesion molecules - have the greatest ability to distinguish among morphologically distinct mouse cortical cell populations (51). Our previous work also suggests that differential expression of ion channel mRNAs in STG cells may give rise to their distinct firing properties (52–54). We therefore examined our scRNA-seq data for expression of ion channel mRNAs. Overall, while the sequencing captured most of the known voltage-gated channel subtypes known in *C. borealis*, raw counts were very low (Fig. 4). Therefore, we decided to use a qRT-PCR approach to directly test the hypothesis that channels and transmitter receptors are effective genes of interest to differentiate known neuron subtypes.

**Figure 4.**
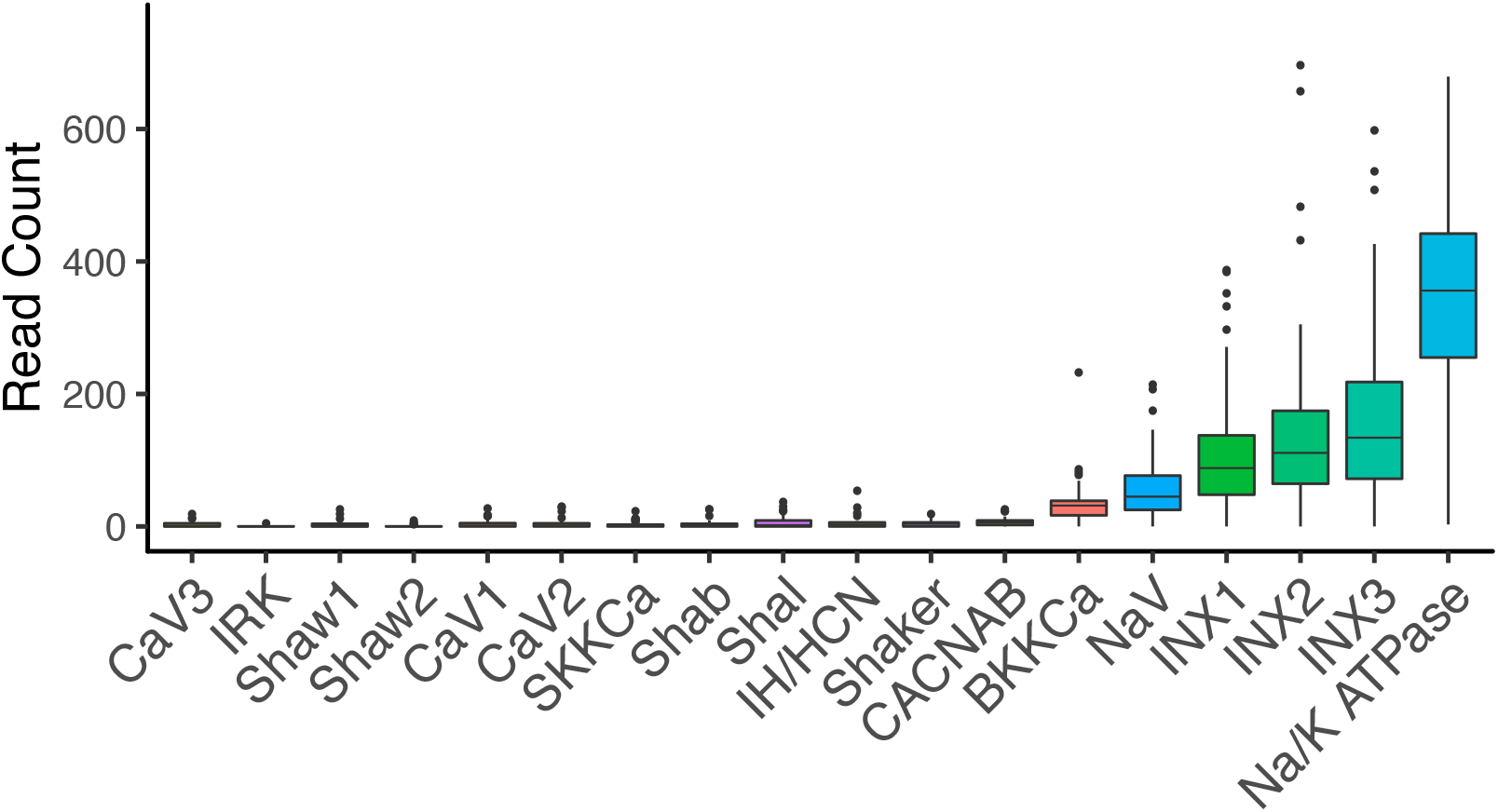
Count numbers for selected voltage-gated ion channels from the RNAseq data. The median counts for each of the voltage-gated channels used in the RT-PCR analysis was generated by pooling cell type. Innexins and the Na+/K+ ATPase are used as a reference of more highly abundant gene products.

To examine the molecular profile of individual identified neurons with qRT-PCR, we targeted the following transcripts: ion channels, receptors, gap junction innexins, and neurotransmitter-related transcripts. These cellular components are responsible for giving neurons much of their unique electrophysiological outputs. As such, we predicted that correspondingly unique expression patterns for this gene set would be present in each neuron type. Using multiplex qRT-PCR, we measured the absolute copy number of 65 genes of interest (see Table S7) from 124 individual STG neurons of 11 different types (10 STG neuron types: PD, LPG (Lateral Posterior Gastric), VD, GM, LP, PY (Pyloric), IC (Inferior Cardiac), LG Lateral Gastric), MG (Median Gastric), DG (Dorsal Gastric) and the Large Cell (LC) motor neurons from the cardiac ganglion (N = 10-15 per type). We then used various methods of unsupervised clustering to generate the “best” clustering of these cells based on *a priori* known cell type. This included substituting any missing values in the qRT-PCR data set via median interpolation.

We then used k-means, unsupervised hierarchical, and SNN-Cliq clustering to generate unbiased clustering analyses based on expression of these genes of interest. Initial interrogation focused on data transformations with a fixed hierarchical clustering scheme (Ward’s D2, Correlation dissimilarity matrix as for the scRNA-seq analysis). Unscaled data as well as centered and data scaled data by gene resulted in different hierarchical clustering patterns. Using unscaled data, hierarchical clustering performed rather poorly in terms of generating distinct clusters that match known cell identity. Performance – as assessed by Jaccard Index – was improved by scaling data across genes, generating 8 distinct nodes with high bootstrap support in hierarchical clustering that capture some of the features of known cell identity (LC, IC, LG, LPG, VD, GM, LP, PD; Fig. 5A). However, multiple cell types fall into clusters that either do not show any separation by neuron identity (DG, MG, PY) or show no bootstrap support based on hierarchical clustering (AU p-value = 0).

**Figure 5.**
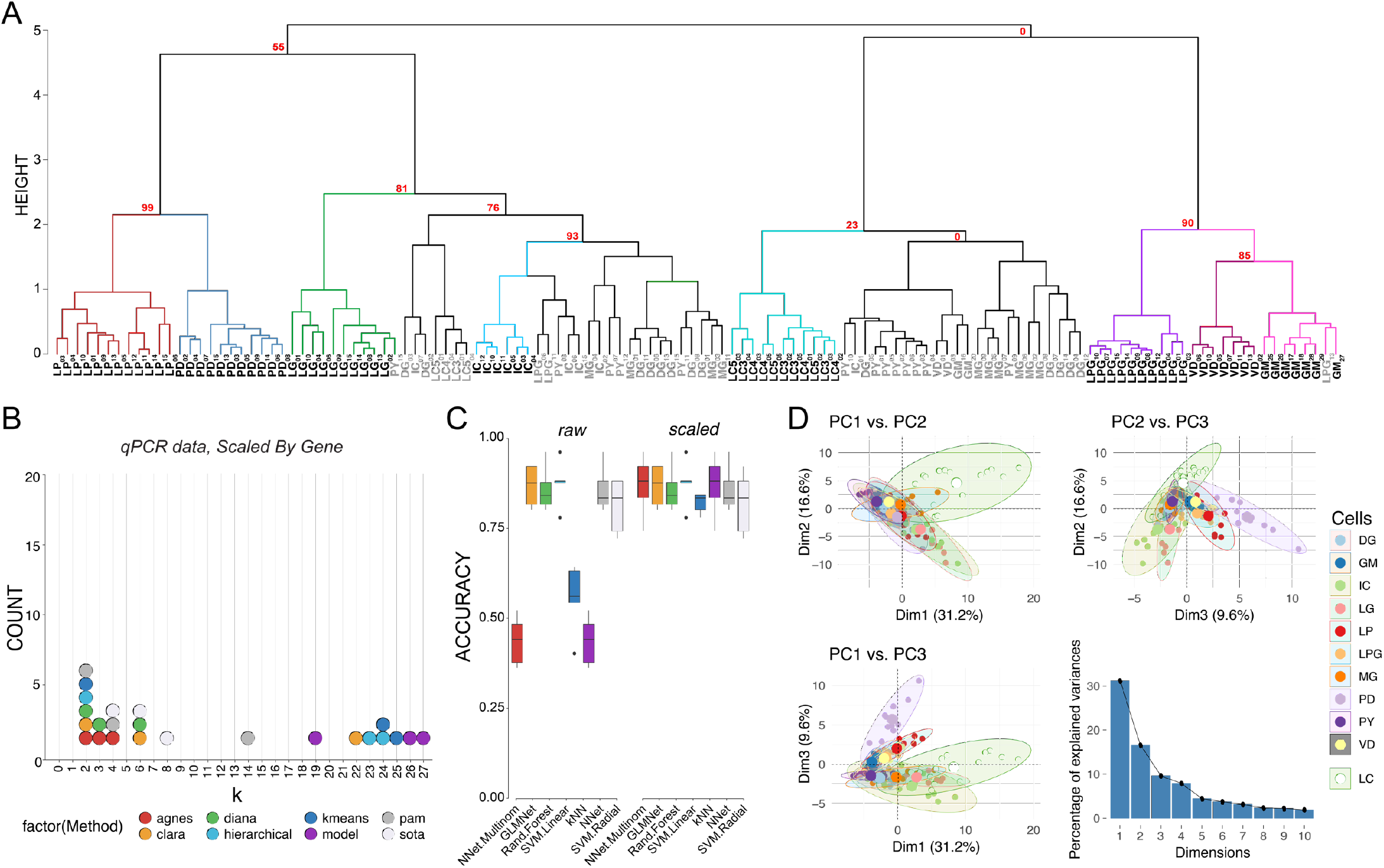
Post-hoc recapitulation of cell identity via qRT-PCR expression with hierarchical clustering and sML algorithms. **A)** Hierarchical clustering of cell type with correlation as the distance metric, and Ward.D2 as the clustering method for data centered and scaled across genes. Approximately Unbiased (AU) p-values for a given node are noted in red. Each node that has >80% support by AU p-value is color coded, and cell types that form a largely coherent group are noted in bold. Cells that do not appear to cluster by type are noted in gray. Cells are identified by type and a subscript that denotes a unique sample identifier. **B)** Dotplot of the top 3 predicted number of clusters based on 8 different prediction algorithms. None of these methods correctly predicted 11 distinct clusters that would represent the 11 different cell types in this assay. **C)** Accuracy of cell type prediction using 8 different methods of sML for each of the data sets. Box and whisker plots show efficacy of each method across five cross-validation folds. **D)** PCA for qRT-PCR data. Pairwise comparisons of PC1, PC2, and PC3 are shown in each panel as in Figure 3. PC1 accounted for 31.2% of the variance, PC2 accounted for 16.6%, and PC3 accounted for 9.6% of the total variance across samples. A scree plot shows the amount of variance explained by PCs 1-10.

We sought to determine the upper bound for clustering performance with our dataset. If the known anatomical and physiological cell identity is reflected in the ion channel and receptor mRNA profile of STG neurons, then we hypothesize that clustering analyses performed on these mRNA data will yield 11 distinct clusters for our dataset. To determine the feasibility of clustering to sort cell types we tested 107 clusterings (varying clustering methods, distance metrics, and neighbors considered) for each data set. Each clustering was compared against the known cell identities with the Jaccard Index which ranges from 0 to 1 where 1 is perfect correspondence between clusterings – in this case the clustering and cell identity. The best performing combination was data scaled by target and processed using Ward’s D2 Hierarchical clustering with a correlation distance matrix (Jaccard = 0.636). By contrast the next best clusterings, Ward’s D on raw counts using Canberra distance and Ward’s D on data scaled by cell using Manhattan distance only achieved Jaccard indices of 0.495 and 0.487 respectively. The three least performant methods were Median hierarchical clustering with Canberra distance (0.084), hierarchical centroid clustering with Manhattan distance (0.859), and SNN-Cliq clustering with Binary distance and 9 neighbors (0.859). Examining the best performing clustering reveals that LP, PD, LG, IC, DG, LC, PY, GM, LPG, and VD separate fairly well.

Given that an *a priori* known number of cell types represented in a sample is rare, we tested whether we would have arrived at the correct number of cell types in our sample had we been blind to their identity. We used the best performing transformations from the clustering analysis, i.e. data centered and scaled by gene and a correlation dissimilarity matrix, and 8 cluster determination indices provided by the optCluster package (49). We allowed a minimum of 2 and a maximum of 32 clusters for this and later cluster determination analyses. The mode of the top 3 predicted k values for 8 different methods of cluster estimation was 2 (6 indices), followed by 4 (the expected number of clusters) and 6 (3 indices each) (see Fig. 5B). If a researcher were using any one of these, or a majority vote of several, the chance they would conclude the correct number of 11 clusters are present would be vanishingly low.

We repeated our sML analyses on the qRT-PCR data to examine the “best case scenario” performance for clustering analyses. Performance varied substantially between algorithms (e.g. NN achieved a mean accuracy of 43.5% whereas SVML produced a mean accuracy of 87.5%) and was affected by whether the data was centered and scaled (e.g. NN improved by 43.5%, SVML did not improve) (Fig. 5C). The highest mean accuracy we achieved was 87.5% (SVML, either with or without scaling). We considered a principal component transformation as well, but improved the maximum mean accuracy little (NN, 87.9%) and worsened the previously most performant methods (SVML decreased from 87.5% to 66.5%, unscaled and 67.4%, scaled). Although neither produces the highest mean accuracy, RF (87.2%-83.2%), GLM (86.6%-79.2%), and LDA (81.9%-77.7%) performed consistently across transformations, but clearly not equally well. Overall, the top performing accuracy methods involved centering and scaling the data across genes, and yielded similar efficacies across algorithms (Fig. 5C).

Finally, we repeated the Principal Component Analysis (PCA) to determine if the variance seen among transcript abundances can be used to separate these 11 cell types into discrete clusters. The first two principal components (PC1 and PC2) generated from the qRT-PCR data accounted for 31.2% and 16.6% of the variance, respectively (Fig. 5D). PC3 accounted for 9.6% of the variance across samples. The top 10 mRNAs contributing to each of these PCs is are listed in Table S1. We generated pairwise plots of all three PCs in attempts to visualize separation of samples into distinct cell types. The most consistent result across all comparisons was that LC neurons from the cardiac ganglion formed a cluster that had less overlap with STG neurons than STG neurons did with each other, particularly in the dimension of PC1 vs. PC2 (Fig. 5D). Visualization of PC1 vs. PC3 and PC2 vs. PC3 also give some indication that even with these target genes of interest we are able to resolve some separation of these groups (Fig. 5D). However, without such extensive *a priori* knowledge about cell type overall it is difficult to see how PCA would be effective in separating these 11 cell types based on the expression data at hand.

### Comparison of qRT-PCR and RNA-seq Results

To ensure that the RNA-seq and qRT-PCR data were producing comparable expression results, we identified 4 different transcripts that were represented both in the DE data set from the RNA-seq and the qRT-PCR data set for the four cell types used in RNA-seq (PD, LP, GM, VD). Overall, there is very strong agreement in expression patterns for all four genes (Fig. 6A), adding confidence to the quality of both data sets with respect to capturing native expression patterns. However, we then extracted the RNA-seq expression data for all 65 of the transcripts used in the qRT-PCR data set. When we performed hierarchical clustering analysis and PCA using these 65 channel and receptor transcripts, the qRT-PCR clusters with nearly 100% success (with the exception of 2 GM neurons) into nodes that contain the 4 known distinct cell types, while the RNA-seq dataset using the same transcripts fails to generate coherent cell type clusters (Fig. 6B; 6C). As we examined this further, we realized that the four transcripts in Figure 6A (*ChAT, vAChT, NMDA2B, KCNK1*) represent somewhat higher abundance transcripts that were differentially expressed and showed consistent patterns between qPCR and RNA-seq methods. Other highly expressed transcript types were not differentially expressed (e.g. *NaV, INX1-3*), and therefore do not contribute strongly to distinguishing cell identity. Conversely, many of the other transcripts in the qRT-PCR data set that were distinct across cell types had very low levels of detected expression in the RNA-seq data set (Fig. 4).

**Figure 6.**
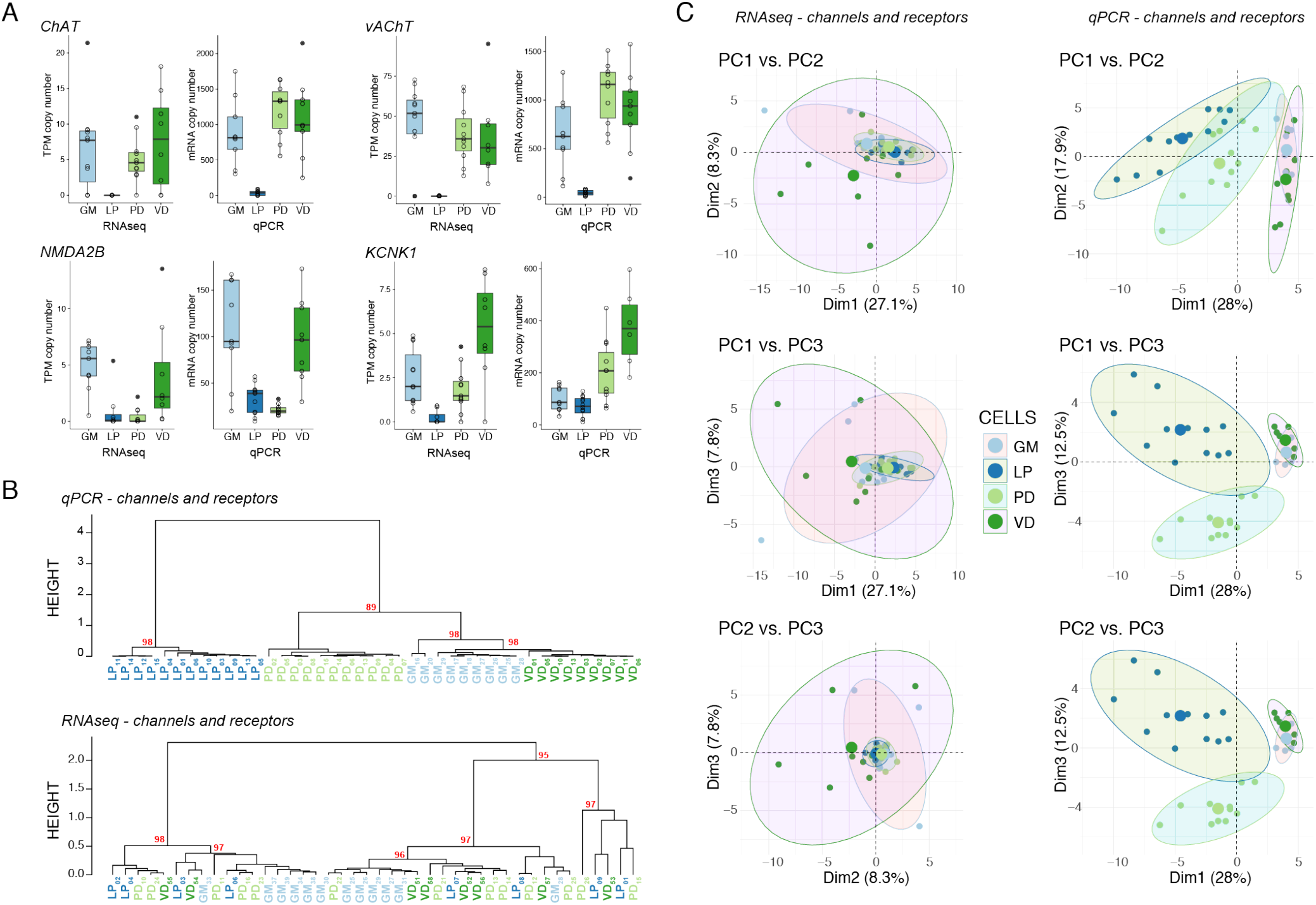
Comparison of expression levels and clustering between qRT-PCR and RNAseq data. **A)** Expression levels of 4 different genes (Choline Acetyltransferase: *ChAT*, Vesicular Acetylcholine Transporter: *vAChT*, NMDA Receptor Subtype 2B: *NMDA2B*, and K^+^ Two-Pore-Domain Channel Subfamily K Member 1: *KCNK1*) between the RNAseq and qRT-PCR data sets. Data shown are medians, quartiles and each individual value from a given animal. Each individual data point is also represented as open circles. RNAseq data are presented as Transcripts Per Kilobase Million (TPM) while qRT-PCR data as absolute copy number per cell. **B)** Hierarchical clustering comparison between qRT-PCR (*top*) and RNAseq (*bottom*) for the same 65 genes represented in the genes of interest pool shown in Figure 1. Each cell type is color coded, and nodes are labeled with AU-values as in previous figures. **C)** PCA for scRNA-seq versus qRT-PCR channel and receptor data. Pairwise comparisons of PC1, PC2, and PC3 are shown in each panel as in Figure 3.

## DISCUSSION

Many projects currently attempting to describe neuronal cell types begin with the acquisition of molecular profiles from populations of unidentified neurons (25, 35, 55). Our results demonstrate the strengths and limitations of both unsupervised and supervised methods that rely solely on a molecular profile to recapitulate neuron identity. We accomplish this by working “backwards” from an unambiguously known cell identity in a system with a rich history of single-cell neurophysiological characterization, the crustacean stomatogastric ganglion. Our analysis clearly demonstrates that even with the most complete *a priori* knowledge of cell type in the analysis, there are limitations to determining cell identity through mRNA expression profiles alone. However, our analyses add to compelling supporting evidence that the molecular profile can partially indicate identity, particularly once supervised methods incorporating known cell identification are employed.

There is increasing evidence that classes of genes may differentiate cell types. For example, genes underlying synaptic transmission machinery were critical for separating mouse cortical GABAergic neurons into different types (56). Sets of genes that are regulated together that can be thought of as a “gene batteries” have also been shown to be indicative of cell type. One well-studied example of this can be found in *C. elegans*, wherein there is expression of neuron-type-specific combinations of transcription factors (57). Most recently, three classes of neuronal effector genes - ion channels, receptors and cell adhesion molecules – were determined to have the greatest ability to distinguish among genetically- and anatomically-defined mouse cortical cell populations (51). Consistent with this work, our GO analysis of the 2000 most variable contigs in our scRNA-seq data set (H2K) revealed that the top 5 Biological Process terms that were significantly enriched included “regulation of short-term neuronal synaptic plasticity,” “substrate adhesion-dependent cell spreading,” and “clathrin-dependent synaptic vesicle endocytosis.” Specifically, our differentially expressed contigs dataset (DE0.2) revealed Molecular Function enrichment for terms related to transmitter identity (“choline:sodium symporter activity” and “acetylcholine transmembrane transporter activity” among others), specifically identified two GABA receptor function terms (“GABA-gated chloride ion channel activity” and “GABA-A receptor activity”) and also included “cell-cell adhesion mediator activity.” Finally, our entire qRT-PCR experiment focused on the expression of ion channels, receptors, gap junction innexins, and neurotransmitter-related transcripts. While these 65 genes were not sufficient for classifying cells into known types, this modest number of transcripts discriminated neuron types fairly well. Thus, categorical families of neuronally expressed genes may yield the most useful data for subdividing neurons into distinct classes or subtypes.

Retinal ganglion cells of mice show spatial patterning in which cells of the same type are distributed with exclusionary zones around them where no other cells of that type are found, while cells of different types do not exhibit spatial patterning and are more randomly distributed (58). Molecular classification of neurons in *C. elegans* found that anatomically distinct neurons have correspondingly distinct molecular profiles >90% of the time (59). However, 146 distinct molecular profiles were identified from the 118 anatomically distinct neuron classes, indicating the potential for molecular sub-classification. This classification relied on hierarchical clustering that was carried out solely on identified reporter genes (most prominently transcription factors and GPCR-type sensory receptors) known to be differentially expressed across the 302 neurons of *C. elegans* from Wormbase.org (60) and not whole transcriptome molecular profiles. It is reassuring that the expression of a wide variety of reporter genes known to be differentially expressed across a population of neurons can recapitulate cell identity. But, this relies on having an established definition of neuron type – in this case anatomical – to constrain hierarchical clustering, as differential expression analysis can only be carried out by assigning samples to different populations. Our results are consistent with these findings, in that clustering is most reliable when differentially expressed targets are present. Yet our data also demonstrate that without separating cell types *a priori by* such additional criteria, molecular cell classification can generate unreliable results, particularly with neurons that belong to the same network.

What are the sources of variability that could mask molecular identification of neuronal identity? Most common high-throughput molecular profiling techniques require destructive sampling to acquire mRNA abundances, which generates only a snapshot of the profile at a single point in time. Gene expression has stochastic characteristics (61, 62); transcription takes place not continually, but in bursts of expression (63) (reviewed in (64)); and steady-state mRNA abundances are the result of rates of expression, but also degradation and mRNA stability (65). Single cell transcriptomes can be altered biologically as a consequence of activity (66), injury (67), long-term memory formation (33), differentiation (68), and aging (23, 69), as well as being affected by technical noise (70).Cells also belong to different transcriptional states under certain conditions, with the major distinction between a cell type and cell state being that state is a reversible condition, where type is more constant and includes neuronal states (71). Neuron types exist in a continuum, exhibiting variation in expression patterns within defined cell types, increasing difficulty in discreetly drawing the cutoff of one type from another (72). Thus, the assertion that a given neuron has a single transcriptomic profile is an oversimplification and simply represents a moment in time in the life of a given cell.

The present study also has limitations. The expression of the focal gene set of ion channels, receptors, gap junction innexins, and neurotransmitter-related transcripts examined in this study ultimately discriminated neuron types fairly well, using supervised methods taking into account known neuron identity. This same gene set did not perform well in the same cell types using RNA-seq (Fig. 6), where a lack of low-abundance transcripts (such as transcription factors and ion channels) may have prevented us from robustly identifying cell-type-specific expression patterns; thus, depth of sequencing is always an ambiguity in every RNA-seq study (73). Furthermore, while we sampled the mRNAt ranscriptome of individual neurons, we have not measured other gene products that could drive unique identity, including non-coding RNA species such as miRNA and lncRNA (74). Epigenetic modifications have also been implicated in neuronal cell identity (75), which were not considered in this study. Further, there are numerous other methods and statistical analyses being applied to molecular profiles to distinguish cell type. We focus on the more commonly employed analyses (PCA, hierarchical clustering, machine learning algorithms) in the literature. Finally, although we are confident in our ability to identify and harvest the targeted neuron types, we cannot rule out the possibility of an occasional misidentified or wrongly isolated cell, as well as the potential presence of adherent support cells.

This study reveals the inherent circularity of the problem facing researchers using transcriptome profiling to identify cell types: molecular profiling is most effective when cells are separated into distinct types *a priori*, yet this is often not possible in many systems. So then how can we most effectively use molecular profiling on unknown populations of cells? The clear answer is to provide as much multimodal data as possible in the analysis. Here, the additional data were an *a priori* separation into cell type based on electrophysiological output, synaptic connectivity, axonal projection, and muscle innervation target (76). While it has been more difficult to achieve multimodal data integration in systems such as cortex, the approach is gaining traction and proving effective. For example, supervised clustering methods proved superior to unsupervised algorithms in separating pyramidal neurons from interneurons in the mouse neocortex based on morphological phenotypes (77). Genetically- and anatomically-defined cell populations in the mouse cortex have revealed much finer resolution and confidence in molecular profiling (51). Much like a circuit’s connectome alone is insufficient to predict network output and function (78), so too the transcriptome alone is insufficient to generate a definitive cell type. Yet it also is clear that transcriptome profiling provides valuable insight into understanding the functional role of individual neurons and neuron types in a network.

## METHODS

### Tissue collection and RNA preparation

Adult Jonah Crabs*, Cancer borealis*, were purchased from the Fresh Lobster Company (Gloucester, Massachusetts, USA) and Commercial Lobster (Boston, MA). Animals were kept in filtered artificial seawater tanks chilled at 10–13°C on a 12/12 light:dark cycle. Prior to dissection, crabs were put on ice for 30 minutes to induce anesthetization. The complete stomatogastric nervous system (STNS) was dissected and pinned out in a dish coated in Sylgard (Dow Corning) with chilled (12°C) physiological saline (composition in mM/l: 440.0 NaCl, 20.0 MgCl2, 13.0 CaCl2, 11.0 KCl, 11.2 Trizma base, and 5.1 maleic acid pH = 7.4 at 23 °C. in RNase-free water). Following desheathing of the stomatogastric ganglion (STG), neurons were identified by simultaneous intra- and extracellular recordings (79, 80). Ten neuron types identified in the STG of *C borealis* were targeted for this study: PD (pyloric dilator), LPG (lateral posterior gastric), LP (lateral pyloric), IC (inferior cardiac), LG (lateral gastric), MG (medial gastric), GM, (gastric mill), PY (pyloric), VD (ventricular dilator), and DG (dorsal gastric). Identified neurons were extracted as previously described (81). Briefly, a Vaseline well was constructed around the ganglion, in which ∼2.5 mg/ml protease (Sigma – P6911, St. Louis, MO) was added to disrupt connective tissue and loosen adherent support cells during a 10-15 minute incubation. The well was then thoroughly washed with fresh physiological saline to halt further enzymatic activity and remove any loose support or connective cells, and a 70% solution of chilled ethylene glycol in saline was added to the well. The saline outside the well was replaced with distilled water, and the entire dish was frozen at −20°C for 30 minutes. This kept the STG neurons cold during the removal of identified neurons. Due to the large size of *C. borealis* STG neuronal somata (50-150 µM in diameter) (82), fine forceps were used to manually remove each neuron. Identified neurons (Fig. 1) were immediately placed in a cryogenic microcentrifuge tube containing 400 µL lysis buffer (Zymo Research) and stored at −80°C until RNA extraction. Total RNA was extracted using the Quick-RNA MicroPrep kit (Zymo Research) per the manufacturer’s protocol.

### Library Preparation and Single-Cell RNA-Seq

Library construction and RNA sequencing services were carried out by the University of Texas at Austin Genomic Sequencing and Analysis Facility (Austin, TX, USA). Extracted single cell RNA from identified neurons from the STG was used to generate cDNA libraries using TruSeq Stranded mRNA Library Prep Kit (Illumina, San Diego, California, USA). Libraries were sequenced in a paired-end 150 bp (2×150bp) configuration on the NextSeq 500 Illumina platform (Illumina, San Diego, CA, USA). Raw reads were processed and analyzed on the Stampede Cluster at the Texas Advanced Computing Facility (TACC). Read quality was checked using the program FASTQC. Low quality reads and adapter sequences were removed using the program Cutadapt (Martin, 2011). The forty identified neurons used in this study all had at least 4 million uniquely mapped reads per sample, comprising 11 PD, 11 GM, 8 LP, and 8 VD cell types. These sequencing reads are deposited in the National Center for Biotechnology Information BioProject archive (PRJNA524309) with the following identifiers: BioSample: SAMN11022125; Sample name: STG Neurons; SRA: SRS4411333.

### Mapping and Differential Expression

The software package Kallisto (83) (v0.43.1) was used in the quantification of RNA-seq abundances through the generation of pseudo-alignments of paired-end fastq files to the *C. borealis* annotated nervous system transcriptome (47). Bootstrapping of the quantification was performed iteratively for 100 rounds. Resulting counts were normalized through the transcripts per kilobase million (TPM) method. Differential expression analysis was carried out using the software package Sleuth (84) (v0.30.0) using TPM normalized counts for each cell type.

### Gene Ontology Enrichment Analysis

Since *C. borealis* lacks a well-curated reference genome, *GO* terms were assigned to the *C. borealis* transcriptome based on best BLASTX hits through reciprocal queries between crab sequence and the *Drosophila melanogaster* NCBI RefSeq database (Release 93). BLAST annotation was carried out based then on Drosophila protein sequence using the BLAST2GO (version 5.1) software suite with the blastx-fast alignment with an E value threshold = 1.0E-3 to generate *D. melanogaster* NCBI Gene IDs associated with each *C. borealis* contig. This produced 1348 and 252 annotated Gene IDs for the H2K and HVG datasets, respectively. These IDs were used as input for statistical overrepresentation tests using the PANTHER Gene Ontology Classification System (v14.1) with default settings using *D. melanogaster* as the reference species. Molecular Function and Biological Process GO terms were examined for enrichment in our datasets, and results reported reflect False Discovery Rate (FDR) correction except where noted.

### Multiplex Primer and Probe Design

Multiplex primer and probe sequences targeting *C. borealis* genes were generated using the RealTimeDesign™ qPCR Assay Design Software from LGC Biosearch Technologies (Petaluma, CA) for custom assays. Multiplex cassettes were designed as a unit to ensure minimal interference in simultaneous qPCR reactions. Probe fluorophore/quencher pairs used in this study are as follows: FAM-BHQ1, CAL Fluor Gold 540-BHQ1, CAL Fluor Red 610-BHQ2, Quasar 670-BHQ2 and Quasar 705-BHQ2. Forward and reverse primer pair, as well as associated probe, sequences can be found in Table S7.

### cDNA synthesis and pre-amplification

Following RNA extraction, individual neuron RNA samples were reverse transcribed into cDNA using qScript cDNA SuperMix (QuantaBio, Beverly, MA, USA) primed with random hexamers and oligo-dT per the manufacturer’s protocol in 20 µL reactions. Half of each resulting cDNA pool (10 µL) was pre-amplified using PerfeCTa PreAmp Supermix (QuantaBio) with a 14-cycle RT-PCR reaction primed with a pool of target-specific primers (Table S7) in a 20 µL reaction per the manufacturer’s protocol to allow for enough product to carry out 15 multiplex qPCR reactions per individual neuron sample. Amplified and unamplified target abundances were compared to ensure minimal amplification bias in the pre-amplification of samples (Fig. S1).

### Quantitative single-cell RT-PCR

Following preamplification of cDNA, samples were diluted 7.5x in nuclease-free water (150 µL final volume) to allow for the quantification of 73 unique gene products across 15 multiplex assays, each able to measure 4-5 different transcripts (Table S7). Reactions were carried out in triplicate on 96-well plates with 10 µL reactions per well using a CFX96 Touch™ Real-Time PCR Detection System from Bio-Rad (Hercules, CA, USA). Cycling conditions for qPCR reactions were as follows: 95°C for 3 min; 40 cycles of 95°C for 15 sec and 58°C for 1 min. Fluorescent measurements were taken at the end of each cycle. The final concentration of primers in each multiplex qPCR reaction was 2.5 µM and 0.3125 µM for each probe.

To quantify absolute mRNA abundances, standard curves were developed for each RT-qPCR multiplex assay using custom gBlock gene fragments (Integrated DNA Technologies, Coralville, IA, USA). Standard curves were generated using a serial dilution of gBlock gene fragments from 1 × 10^6^ to 1 × 10^1^ copies for each reaction assay and were shown to be linear and reproducible. Copy numbers were calculated using the efficiency and slope generated from the standard curves and accounting for the 14-cycle preamplification and subsequent cDNA dilution described above.

### Statistical Analysis

All statistical analyses were performed using R version 3.5.3 (2019-03-11) -- “Great Truth” (85). We used single cell RNA-seq data to evaluate our methods under expected and near best case scenarios. To this end, we reduced the dimensionality of the data (28,695 contigs) by selecting the 2000 most variable contig and by selecting 922 highly variable contigs selected using the M3Drop implementation of the Brennecke method (48) (i.e. M3Drop:: BrenneckeGetVariableGenes() (86)) assuming a 0.2 false discovery rate. To test performance under ideal conditions we selected those contigs differentially expressed at an alpha of 0.2 or 0.05. We centered and scaled the aforementioned datasets and their progenitors via the caret::predict() and caret::preprocess() functions (87). We also tested dimensionality reduction via PCA. We further used PCA in exploratory data analysis to determine if any of the cell types were visually separable across four subsets of the data (Seq H2K, Seq HVG, Seq DE0.2, and Seq DE0.05).

Next, we performed cluster estimation using the optClust() function of the optCluster package (49). The algorithms used on each dataset varied by whether the data were counts or continuous. Allowed k values ranged from 2-10 (i.e. cells in dataset / 4, rounding up). We selected the top three predicted k values from each algorithm for visualization of the spread of predicted ks.

To assess the performance of unsupervised machine learning methods on our data we tested several clustering algorithms – k-means clustering, hierarchical clustering (using a variety of distance metrics, (euclidean, maximum, manhattan, canberra, binary, minkowski, correlation, uncentered) and clustering methods (ward.D, ward.D2, single, complete, average, mcquitty, median, centroid, ward.D2)), and SNN-Cliq clustering (88). We then selected high performing clustering methods based on the Jaccard index calculated against cell identity. We selected one of the best performing combinations (Ward’s method with correlation as the distance metric) for visualization.

We applied several supervised machine learning methods to evaluate predictive power of expression data in ideal circumstances (i.e. prior knowledge of a given cell type’s molecular identity). Specifically, we tested elastic regression, k-nearest neighbors, linear discriminant analysis, neural network, multinomial neural network, random forest, support vector machine with a radial kernel, and support vector machine with a linear kernel. For each of these models we tested a variety of tuning parameters and selected the most effective parameter set before comparison with other methods. Methods were evaluated by using cross validation (with five folds) to produce the expected accuracy on new data. The same approaches were applied to the single cell RT-qPCR data set, with a few caveats. Given its relatively smaller size, dimensionality reduction was not necessary to overcome technical or practical hurdles. Thus, we tested both the raw and centered and scaled dataset in addition to PCA transformations of the same. We also increased the maximum k allowed in cluster estimation to 32.

## DECLARATIONS

### Availability of Data and Material

All sequence data accession numbers are provided in the manuscript and accompanying tables.

### Authors Contributions

Conceived the study: DJS, EM, HAH. Tissue collection: AJN, AGO, JMS, RMH. Performed qPCR assays: AJN. Design of Primers: AJN, DJS. RNA-seq analysis: BMG, AJN, DRK, DJS, HAH, RMH. Data analysis: AJN, DRK, DJS. Interpretation of results: AJN, DRK, DJS, EM, HAH. Wrote the manuscript: AJN, DRK, DJS, EM, HAH. All authors read and approved the final manuscript.

## Acknowledgements

We would like to thank members of the Schulz and Hofmann labs for helpful discussions. The authors thank the Genomic Sequencing and Analysis Facility at UT Austin for library preparation and sequencing and the bioinformatics consulting team at the UT Austin Center for Computational Biology and Bioinformatics for helpful advice.

## SUPPLEMENTAL FIGURES

**Figure S1.**
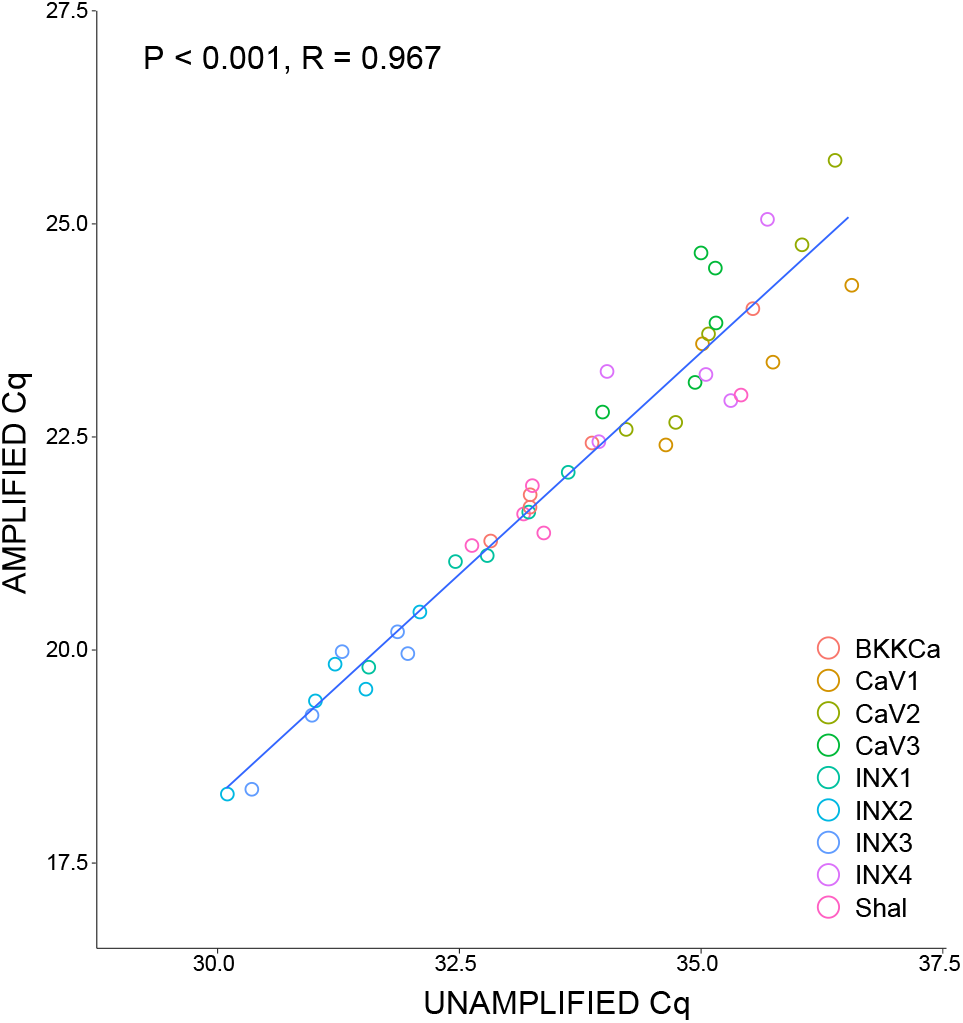
Comparison of expression levels of the same single cell samples (N = 5) before (Unamplified) and after (Amplified) 14 cycles of preamplification of cDNA. Each sample represents a cDNA pool from a single identified neuron, half of which was preamplified and half remained unamplified. Data are shown as quantitation cycle (Cq) values. Statistics shown report values for Pearson’s Correlation test.

**Table S1.** Top ten contributing genes or contigs to PCs1-3 for each dataset.

| **Dataset** | **Rank** | **PC1** | **PC1 value** | **PC2** | **PC2 value** | **PC3** | **PC3 value** |
| --- | --- | --- | --- | --- | --- | --- | --- |
| *qRTPCR* | 1 | LCCH3r | 3.54 | NaV | 6.30 | GluCl | 9.12 |
| *qRTPCR* | 2 | mGABA3 | 3.10 | Shal | 5.65 | GABAB R1 | 8.04 |
| *qRTPCR* | 3 | mGluR5 | 3.06 | NMDA_1A | 5.16 | vAChT | 7.56 |
| *qRTPCR* | 4 | CaV2 | 3.05 | KCNK1 | 4.95 | HisCL | 6.74 |
| *qRTPCR* | 5 | Shab | 3.02 | Shaker | 4.54 | ChAT | 6.57 |
| *qRTPCR* | 6 | KCNH3 | 2.97 | IH | 3.47 | IH | 6.16 |
| *qRTPCR* | 7 | DAR1A | 2.91 | KCNK2 | 3.45 | INX4 | 6.13 |
| *qRTPCR* | 8 | SKKCa | 2.87 | BKKCa | 3.10 | vGluT | 5.29 |
| *qRTPCR* | 9 | His 3r | 2.86 | Dopa 1Br | 3.06 | RDLr | 4.16 |
| *qRTPCR* | 10 | TRP A like | 2.82 | RDLr | 2.93 | CCAPr | 3.11 |
| *seq_h2k* | 1 | c1318 | 0.10 | c4636 | 0.50 | c4191 | 0.59 |
| *seq_h2k* | 2 | c724 | 0.10 | c8463 | 0.50 | c751 | 0.58 |
| *seq_h2k* | 3 | c2022 | 0.10 | c28755 | 0.50 | c8533 | 0.49 |
| *seq_h2k* | 4 | c1834 | 0.10 | c17319 | 0.50 | c953 | 0.46 |
| *seq_h2k* | 5 | c718 | 0.10 | c10220 | 0.50 | c2665 | 0.46 |
| *seq_h2k* | 6 | c2357 | 0.10 | c27163 | 0.49 | c2126 | 0.45 |
| *seq_h2k* | 7 | c196 | 0.10 | c5528 | 0.49 | c2981 | 0.45 |
| *seq_h2k* | 8 | c739 | 0.10 | c10716 | 0.49 | c3881 | 0.42 |
| *seq_h2k* | 9 | c2301 | 0.10 | c9333 | 0.49 | c23433 | 0.41 |
| *seq_h2k* | 10 | c1048 | 0.10 | c13463 | 0.49 | c1647 | 0.40 |
| *seq_hvg* | 1 | c38450 | 0.50 | c18443 | 0.63 | c29394 | 0.80 |
| *seq_hvg* | 2 | c5595 | 0.50 | c17911 | 0.63 | c23916 | 0.80 |
| *seq_hvg* | 3 | c28755 | 0.50 | c13615 | 0.63 | c39794 | 0.80 |
| *seq_hvg* | 4 | c11256 | 0.49 | c16416 | 0.63 | c24360 | 0.80 |
| *seq_hvg* | 5 | c20433 | 0.49 | c17569 | 0.63 | c18403 | 0.79 |
| *seq_hvg* | 6 | c39762 | 0.49 | c17622 | 0.63 | c7694 | 0.79 |
| *seq_hvg* | 7 | c13489 | 0.49 | c18306 | 0.63 | c11984 | 0.79 |
| *seq_hvg* | 8 | c19224 | 0.49 | c19165 | 0.63 | c16991 | 0.79 |
| *seq_hvg* | 9 | c30088 | 0.49 | c19999 | 0.63 | c18899 | 0.79 |
| *seq_hvg* | 10 | c4923 | 0.49 | c22142 | 0.63 | c25542 | 0.79 |
| *seq_DE0.2* | 1 | c5749 | 1.62 | c4517 | 3.20 | c13441 | 5.94 |
| *seq_DE0.2* | 2 | c1898 | 1.44 | c398 | 2.92 | c1058 | 4.27 |
| *seq_DE0.2* | 3 | c23967 | 1.43 | c878 | 2.92 | c31757 | 3.14 |
| *seq_DE0.2* | 4 | c5120 | 1.41 | c4945 | 2.87 | c9248 | 3.05 |
| *seq_DE0.2* | 5 | c1219 | 1.39 | c3559 | 2.69 | c2212 | 3.02 |
| *seq_DE0.2* | 6 | c8871 | 1.37 | c15559 | 2.64 | c4534 | 2.63 |
| *seq_DE0.2* | 7 | c972 | 1.35 | c8507 | 2.50 | c8114 | 2.61 |
| *seq_DE0.2* | 8 | c973 | 1.33 | c1151 | 2.15 | c10145 | 2.57 |
| *seq_DE0.2* | 9 | c12663 | 1.32 | c8323 | 2.06 | c2981 | 2.47 |
| *seq_DE0.2* | 10 | c910 | 1.30 | c1800 | 2.00 | c14660 | 2.35 |
| *seq_DE0.05* | 1 | c21272 | 4.69 | c4517 | 8.85 | c1058 | 13.16 |
| *seq_DE0.05* | 2 | c5716 | 4.21 | c16963 | 7.16 | c13441 | 11.60 |
| *seq_DE0.05* | 3 | c5120 | 4.14 | c3348 | 6.68 | c2212 | 10.64 |
| *seq_DE0.05* | 4 | c3737 | 3.77 | c1151 | 6.05 | c2586 | 7.17 |
| *seq_DE0.05* | 5 | c8114 | 3.72 | c14320 | 5.14 | c5845 | 5.83 |
| *seq_DE0.05* | 6 | c16240 | 3.59 | c5222 | 4.93 | c14320 | 4.87 |
| *seq_DE0.05* | 7 | c2796 | 3.56 | c8323 | 4.57 | c4997 | 4.10 |
| *seq_DE0.05* | 8 | c1713 | 3.43 | c5067 | 4.35 | c14660 | 4.06 |
| *seq_DE0.05* | 9 | c49 | 3.38 | c1324 | 4.12 | c3348 | 4.00 |
| *seq_DE0.05* | 10 | c3716 | 3.23 | c24846 | 4.08 | c4534 | 3.86 |

**Table S2.** Gene Ontology Enrichment analysis of Molecular Function for H2K RNAseq data.

| **GO term: molecular function** | **Fold**<br>**Enrichment** | **FDR**<br>**p-value** |
| --- | --- | --- |
| proton-transporting ATP synthase activity, rotational mechanism (GO:0046933) | 8.08+ | 2.88E-04 |
| clathrin binding (GO:0030276) | 6.82+ | 4.51E-03 |
| ubiquitin conjugating enzyme activity (GO:0061631) | 4.6+ | 1.59E-02 |
| intramolecular oxidoreductase activity (GO:0016860) | 4.32+ | 2.22E-02 |
| structural constituent of ribosome (GO:0003735) | 4.02+ | 4.61E-11 |
| proton-transporting ATPase activity, rotational mechanism (GO:0046961) | 3.84+ | 4.35E-02 |
| structural constituent of cytoskeleton (GO:0005200) | 3.84+ | 4.30E-02 |
| unfolded protein binding (GO:0051082) | 3.8+ | 4.00E-05 |
| heat shock protein binding (GO:0031072) | 3.73+ | 5.01E-02 |
| mRNA 3'-UTR binding (GO:0003730) | 3.73+ | 4.96E-02 |
| cell adhesion molecule binding (GO:0050839) | 3.57+ | 3.89E-02 |
| translation factor activity, RNA binding (GO:0008135) | 3.27+ | 4.57E-03 |
| electron transfer activity (GO:0009055) | 3.07+ | 1.79E-02 |
| GTPase activity (GO:0003924) | 3.07+ | 6.09E-05 |
| GTP binding (GO:0005525) | 2.97+ | 6.78E-05 |
| actin binding (GO:0003779) | 2.95+ | 4.42E-04 |
| kinase binding (GO:0019900) | 2.85+ | 7.78E-03 |
| microtubule binding (GO:0008017) | 2.75+ | 3.98E-03 |
| phospholipid binding (GO:0005543) | 2.56+ | 3.16E-02 |
| calcium ion binding (GO:0005509) | 2.4+ | 1.60E-03 |
| protein-containing complex binding (GO:0044877) | 2.39+ | 7.29E-04 |
| ATP binding (GO:0005524) | 2.31+ | 1.36E-10 |
| protein serine/threonine kinase activity (GO:0004674) | 2.17+ | 1.31E-02 |
| enzyme regulator activity (GO:0030234) | 1.83+ | 1.40E-02 |
| DNA-binding transcription factor activity (GO:0003700) | 0.41- | 1.79E-02 |
| serine-type endopeptidase activity (GO:0004252) | 0.1- | 1.90E-03 |

**Table S3.** Gene Ontology Enrichment analysis of Biological Process for H2K RNAseq data.

| **GO term: biological process** | **Fold Enrichment** | **FDR p-value** |
| --- | --- | --- |
| regulation of short-term neuronal synaptic plasticity (GO:0048172) | 10.96+ | 6.64E-03 |
| positive regulation of neuron remodeling (GO:1904801) | 10.63+ | 7.39E-05 |
| substrate adhesion-dependent cell spreading (GO:0034446) | 10.23+ | 2.68E-03 |
| actin filament polymerization (GO:0030041) | 9.59+ | 9.52E-03 |
| clathrin-dependent synaptic vesicle endocytosis (GO:0150007) | 8.77+ | 3.43E-02 |
| protein N-linked glycosylation via asparagine (GO:0018279) | 8.77+ | 3.42E-02 |
| gluconeogenesis (GO:0006094) | 8.53+ | 1.31E-02 |
| dorsal closure, spreading of leading edge cells (GO:0007395) | 8.53+ | 1.31E-02 |
| morphogenesis of larval imaginal disc epithelium (GO:0016335) | 7.67+ | 4.63E-02 |
| retrograde axonal transport (GO:0008090) | 7.67+ | 1.79E-02 |
| ATP synthesis coupled proton transport (GO:0015986) | 7.03+ | 1.14E-04 |
| vesicle transport along microtubule (GO:0047496) | 6.98+ | 2.40E-02 |
| axonal transport of mitochondrion (GO:0019896) | 6.98+ | 2.39E-02 |
| anterograde axonal transport (GO:0008089) | 6.98+ | 2.39E-02 |
| cellular response to metal ion (GO:0071248) | 6.39+ | 3.07E-02 |
| axonal fasciculation (GO:0007413) | 6.14+ | 3.23E-03 |
| female germ-line stem cell asymmetric division (GO:0048132) | 5.97+ | 7.80E-03 |
| regulation of reactive oxygen species metabolic process (GO:2000377) | 5.9+ | 3.80E-02 |
| actin nucleation (GO:0045010) | 5.48+ | 4.75E-02 |
| positive regulation of photoreceptor cell differentiation (GO:0046534) | 5.48+ | 4.74E-02 |
| sevenless signaling pathway (GO:0045500) | 5.42+ | 2.40E-02 |
| ovarian follicle cell stalk formation (GO:0030713) | 5.42+ | 2.39E-02 |
| synaptic vesicle priming (GO:0016082) | 5.42+ | 2.39E-02 |
| flight behavior (GO:0007629) | 5.42+ | 2.39E-02 |
| positive regulation of endocytosis (GO:0045807) | 5.42+ | 3.15E-04 |
| positive regulation of lipid localization (GO:1905954) | 5.42+ | 2.38E-02 |
| positive regulation of canonical Wnt signaling pathway (GO:0090263) | 5.39+ | 1.58E-04 |
| ribosomal large subunit assembly (GO:0000027) | 5.37+ | 1.19E-02 |
| positive regulation of smoothened signaling pathway (GO:0045880) | 5.31+ | 3.15E-03 |
| positive regulation of protein modification by small protein conjugation or removal (GO:1903322) | 5.12+ | 2.94E-02 |
| glucose homeostasis (GO:0042593) | 5.12+ | 2.40E-04 |
| lumen formation, open tracheal system (GO:0035149) | 5.12+ | 3.83E-03 |
| cellular protein complex disassembly (GO:0043624) | 5.12+ | 7.47E-03 |
| glycolytic process (GO:0006096) | 4.91+ | 8.93E-03 |
| oocyte microtubule cytoskeleton polarization (GO:0008103) | 4.85+ | 3.49E-02 |
| hemocyte migration (GO:0035099) | 4.6+ | 4.17E-02 |
| negative regulation of supramolecular fiber organization (GO:1902904) | 4.55+ | 1.26E-02 |
| maintenance of protein location in cell (GO:0032507) | 4.51+ | 4.05E-03 |
| regulation of R7 cell differentiation (GO:0045676) | 4.39+ | 4.95E-02 |
| cell adhesion mediated by integrin (GO:0033627) | 4.39+ | 4.94E-02 |
| positive regulation of neuromuscular junction development (GO:1904398) | 4.39+ | 4.76E-03 |
| terminal branching, open tracheal system (GO:0007430) | 4.39+ | 1.49E-02 |
| pole plasm oskar mRNA localization (GO:0045451) | 4.32+ | 9.07E-03 |
| positive regulation of organ growth (GO:0046622) | 4.3+ | 3.01E-02 |
| antibiotic metabolic process (GO:0016999) | 4.3+ | 3.00E-02 |
| adherens junction organization (GO:0034332) | 4.26+ | 5.62E-03 |
| epithelial cell migration, open tracheal system (GO:0007427) | 4.26+ | 5.61E-03 |
| regulation of cell-cell adhesion (GO:0022407) | 4.23+ | 1.77E-02 |
| intestinal stem cell homeostasis (GO:0036335) | 4.23+ | 1.77E-02 |
| regulation of filopodium assembly (GO:0051489) | 4.19+ | 1.06E-02 |
| heart morphogenesis (GO:0003007) | 4.15+ | 6.50E-03 |
| negative regulation of autophagy (GO:0010507) | 4.13+ | 3.47E-02 |
| reactive oxygen species metabolic process (GO:0072593) | 4.13+ | 3.45E-02 |
| plasma membrane invagination (GO:0099024) | 4.13+ | 3.45E-02 |
| ATP hydrolysis coupled proton transport (GO:0015991) | 4.04+ | 7.57E-03 |
| olfactory learning (GO:0008355) | 4.04+ | 7.54E-03 |
| positive regulation of translation (GO:0045727) | 4.04+ | 7.50E-03 |
| rhabdomere development (GO:0042052) | 4.02+ | 4.61E-03 |
| synaptic growth at neuromuscular junction (GO:0051124) | 3.95+ | 1.43E-02 |
| protein folding (GO:0006457) | 3.93+ | 6.74E-08 |
| establishment of mitotic spindle localization (GO:0040001) | 3.84+ | 4.66E-02 |
| salivary gland cell autophagic cell death (GO:0035071) | 3.84+ | 4.65E-02 |
| insulin receptor signaling pathway (GO:0008286) | 3.84+ | 4.64E-02 |
| regulation of lipid storage (GO:0010883) | 3.74+ | 1.15E-02 |
| negative regulation of cytoskeleton organization (GO:0051494) | 3.72+ | 3.20E-02 |
| negative regulation of smoothened signaling pathway (GO:0045879) | 3.72+ | 3.20E-02 |
| mitochondrial ATP synthesis coupled electron transport (GO:0042775) | 3.65+ | 1.75E-04 |
| regulation of peptide secretion (GO:0002791) | 3.64+ | 2.21E-02 |
| behavioral response to ethanol (GO:0048149) | 3.63+ | 3.40E-03 |
| regulation of chemotaxis (GO:0050920) | 3.61+ | 3.63E-02 |
| response to unfolded protein (GO:0006986) | 3.61+ | 3.62E-02 |
| positive regulation of cell size (GO:0045793) | 3.57+ | 1.48E-02 |
| tight junction organization (GO:0120193) | 3.54+ | 2.50E-02 |
| apical junction assembly (GO:0043297) | 3.52+ | 1.02E-02 |
| cytosolic transport (GO:0016482) | 3.5+ | 4.39E-03 |
| cytokinetic process (GO:0032506) | 3.49+ | 1.69E-02 |
| regulation of axonogenesis (GO:0050770) | 3.47+ | 3.00E-03 |
| negative regulation of protein phosphorylation (GO:0001933) | 3.44+ | 2.05E-03 |
| positive regulation of cell morphogenesis involved in differentiation (GO:0010770) | 3.41+ | 4.69E-02 |
| establishment or maintenance of apical/basal cell polarity (GO:0035088) | 3.37+ | 3.20E-02 |
| regulation of multicellular organism growth (GO:0040014) | 3.34+ | 2.20E-02 |
| morphogenesis of follicular epithelium (GO:0016333) | 3.31+ | 1.47E-02 |
| translational initiation (GO:0006413) | 3.25+ | 1.65E-02 |
| regulation of protein stability (GO:0031647) | 3.25+ | 1.65E-02 |
| neuromuscular synaptic transmission (GO:0007274) | 3.23+ | 1.13E-02 |
| autophagy (GO:0006914) | 3.23+ | 4.15E-04 |
| imaginal disc-derived wing margin morphogenesis (GO:0008587) | 3.19+ | 1.86E-02 |
| mitotic cytokinesis (GO:0000281) | 3.16+ | 6.07E-03 |
| long-term memory (GO:0007616) | 3.15+ | 4.15E-03 |
| germ-line stem cell population maintenance (GO:0030718) | 3.15+ | 2.84E-03 |
| asymmetric neuroblast division (GO:0055059) | 3.14+ | 4.50E-02 |
| cell redox homeostasis (GO:0045454) | 3.13+ | 3.09E-02 |
| response to growth factor (GO:0070848) | 3.13+ | 3.08E-02 |
| synaptic target recognition (GO:0008039) | 3.13+ | 3.07E-02 |
| amino acid transport (GO:0006865) | 3.01+ | 3.76E-02 |
| axon guidance (GO:0007411) | 2.94+ | 1.77E-08 |
| positive regulation of locomotion (GO:0040017) | 2.63+ | 4.08E-02 |
| negative regulation of neurogenesis (GO:0050768) | 2.63+ | 3.09E-02 |

**Table S4.** Gene Ontology Enrichment analysis of Molecular Function for HVG RNAseq data.

| **GO term: molecular function** | **Fold Enrichment** | **FDR p-value** |
| --- | --- | --- |
| ATP binding (GO:0005524) | 3.1+ | 1.72E-03 |
| transferase activity (GO:0016740) | 2+ | 2.38E-02 |

**Table S5.** Gene Ontology Enrichment analysis of Biological Process for HVG RNAseq data.

| **GO term: molecular function** | **Fold Enrichment** | **FDR p-value** |
| --- | --- | --- |
| regulation of protein localization to plasma membrane (GO:1903076) | 35.33+ | 4.79E-02 |
| chromatin silencing (GO:0006342) | 7.69+ | 3.12E-02 |
| nucleic acid metabolic process (GO:0090304) | 2.17+ | 3.74E-02 |
| macromolecule modification (GO:0043412) | 2.08+ | 3.91E-02 |
| cellular macromolecule metabolic process (GO:0044260) | 1.84+ | 2.22E-02 |

**Table S6.**
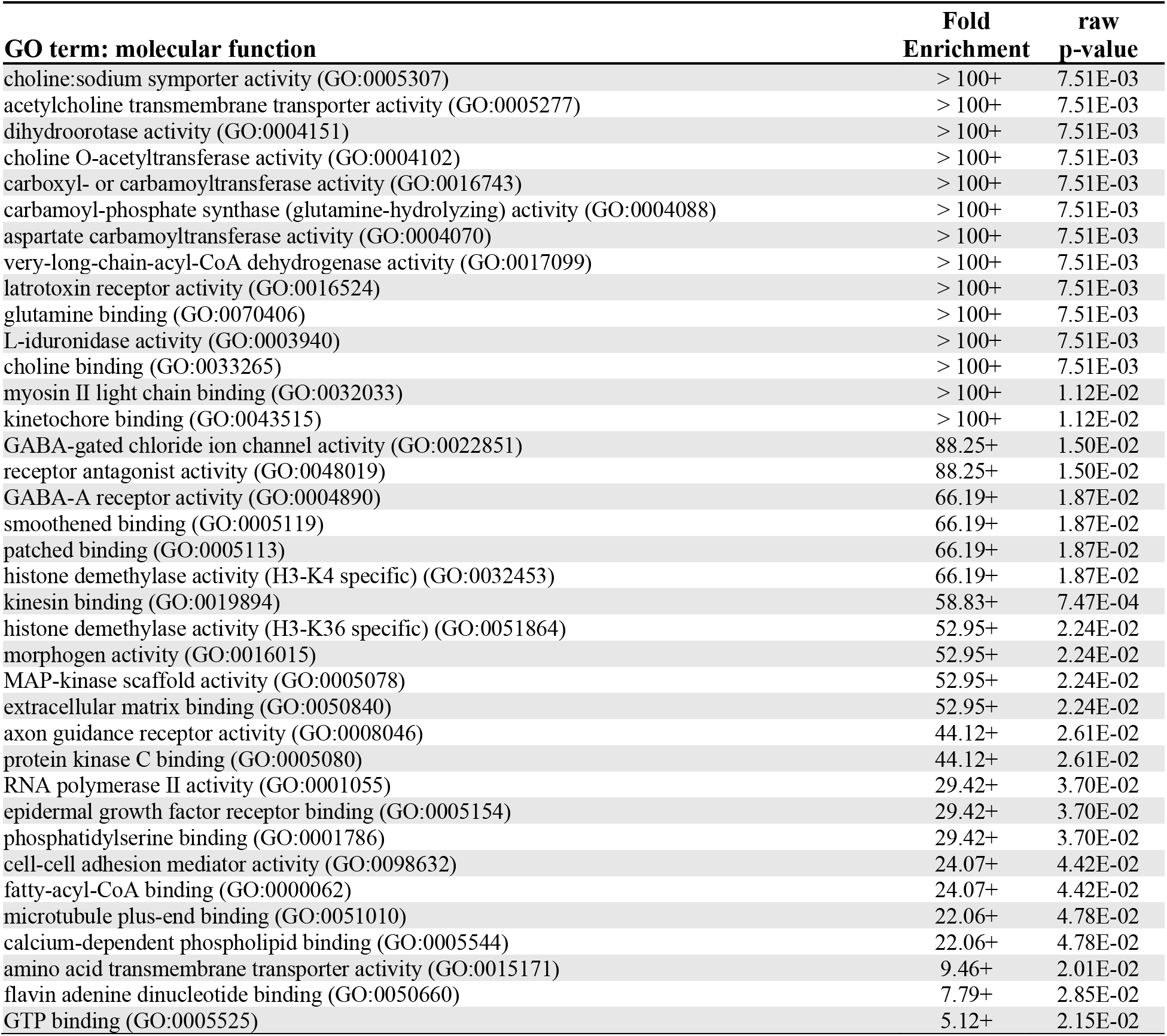
Gene Ontology Enrichment analysis of Molecular Function for DE0.2 RNAseq data.

**Table S7.**
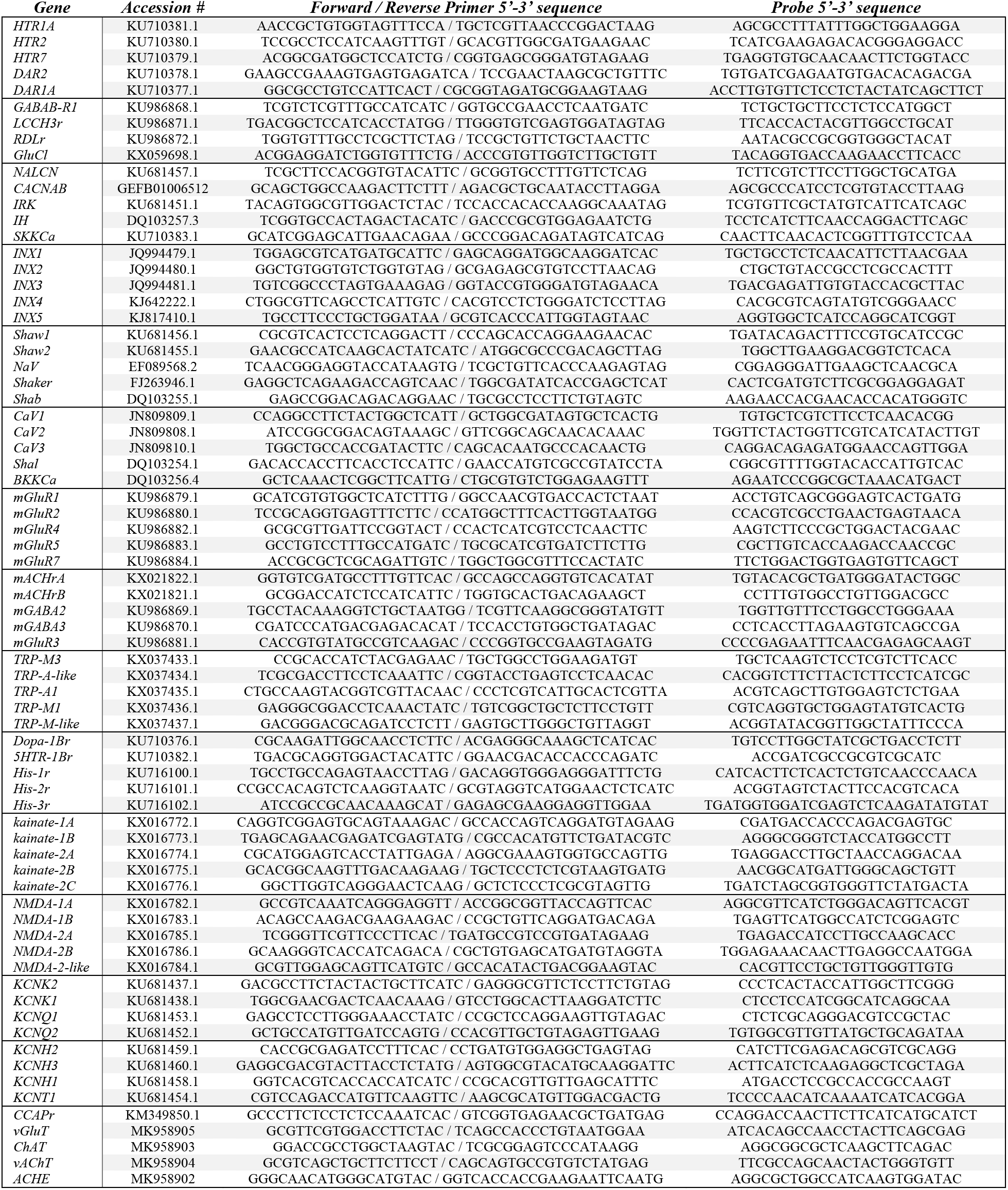
Target primer and probe sequences for qRT-PCR Multiplex assays. Each box represents a group of four to five genes that were combined into a single multiplex reaction.

